# Chicoric acid prevents neurodegeneration via microbiota-gut-brain axis in a mouse Parkinson’s disease model

**DOI:** 10.1101/2021.05.03.442390

**Authors:** Ning Wang, Xin-Yue Pan, Hong-Kang Zhu, Ya-Hui Guo, He Qian

**Affiliations:** School of Food Science and Technology, Jiangnan University, Wuxi214000, China

**Keywords:** Parkinson’s Disease, chicoric acid, microbiota-gut-brain axis

## Abstract

It is determined that microbiota-gut-brain axis is involved in the pathogenesis of Parkinson’s Disease (PD) and may be the potential target for developing the therapeutic treatments for PD. Chicoric acid (CA) is a kind of natural polyphenolic acid compounds which are recognized as promising agents against neurodegenerative disease. Here, we investigated the influence of CA on microbiota-gut-brain axis in 1-methyl-4-phenyl-1, 2, 3, 6-tetrahydropyridine (MPTP) mice. The results demonstrated that oral pretreatments of CA significantly prevented the MPTP-indued motor disorders, death of nigrostriatal dopaminergic neurons and reduction in striatal neurotrophins. Sequencing results of 16S rRNA indicated the microbial dysbiosis occurred in MPTP mice, whereas CA exerted a remarkable impact on microbial diversity and microbiota compositions, as well as SCFAs production. Besides, CA pretreatment alleviated gliosis-mediated neuroinflammation and gut inflammation by suppressing TLR4/MyD88/NF-*κ*B signaling cascade, along with preventing the colonic hyperpermeability. To conclude, CA prevented MPTP-induced neuroinflammation and neuronal degenerative processes through signaling multiple pathways of microbiota-gut-brain axis.

## Introduction

Parkinson’s disease (PD) is the second most prevalent neurodegenerative disorder, characterized by progressive degeneration of dopaminergic neurons in the substantia nigra pars compacta (SNpc). The disease is manifested in various progressive motor deficits, including resting tremor, bradykinesia, and muscle stiffness. In addition, as the pathology extends to cortical and subcortical non-dopaminergic degeneration, non-motor dysfunctions such as depression, delusions and cognitive impairments occur in PD (Jankovic, 2008; Perlmutter et al., 2020). So far, there is no effective clinical treatments for PD, thus it is urgent to explore new therapies and/or targets.

It was noted that multiple gastrointestinal (GI) abnormalities including weight loss, dysphagia, and constipation always precede motor-symptoms by many years as common PD manifestations(Verbaan et al., 2007), suggesting the linkage between the PD pathogenesis and GI dysfunctions of involving gut microbial imbalance (Kim et al., 2015). Recent reported works demonstrated the crucial role of gut microbial dysbiosis induced imbalance of the gut-microbiome-brain axis in pathogenesis and development of PD (Baizabal-Carvallo et al., 2020). To date, evidences are still being accumulated and increasing works are still under conducting to elucidate the interactions between microbiota-gut-brain axis and PD pathology.

Chicoric acid (CA), a natural polyphenolic acid from chicory and Echinacea purpurea, is described as a potential agent promoting health depending on its anti-inflammatory, antioxidative, immunoregulatory and antiviral activities, it has been used as medication or nutraceutical against diabetes, epilepsy, eye diseases, digestive disorders and flu (Ahn et al., 2014; Street et al., 2013). CA was found to be a polyphenol compounds of which most have an effect on modulation of the gut microbial population (Cardona et al., 2013). Of note, polyphenols were indicated to be highly advantageous to counteract inflammation and oxidative stress in the brain owing to their abilities interplaying with brain cells directly, as well as modulating microbiota-gut-brain axis (Serra et al., 2020). Moreover, previous studies suggested that CA could inhibit neuroinflammation in vivo and in vitro by downregulating NF-*κ*B pathway, and exert neuroprotective effects by preventing oxidative stress (Liu et al., 2017; Wang et al., 2017). However, the effect of CA on neuronal survivals in PD is still unknow. Hence, we hypothesized that CA pretreatment could protect dopaminergic neuronal loss by inhibit inflammation through signaling microbiota-gut-brain_Ú_

Accumulating reports have indicated that gut microbiota composition was heavily involved in gut-brain axis of PD. For instance, both mice and patients with PD possessed an enhancement of phylum *proteobacteria*, which concede to several pathogens due to their pro-inflammation properties (Gorecki et al., 2019; Jang et al., 2020). Gerhardt et al. found a decrease in genus *Butyricimona*, a butyrate producer with anti-inflammatory properties. Additionally, phylum *Firmicute*s in PD were reported and confirmed to change the most in previous studies, as an increase of Lactobacillus, *Bifidobacterium, Verrucomicrobiaceae* and *Akkermansia*, and a decrease of *Faecalibacterium, Coprococcus, Blautia, Prevotella* and *Prevotellaceae* were observed(Gerhardt et al., 2018), while Lactobacillus, a beneficial gut microbe synthesizing vitamin B12, was shown decreased in Qian’s works(LeBlanc et al., 2013; Qian et al., 2018), which is also consistent with this findings. Although different results were accomplished in these studies, distinct changes actually occurred in gut microbial composition in PD.

It is well documented that inflammatory processes participate in the neuronal degeneration throughout the initiation and development of PD. Recent studies provided the evidences that neuroinflammation came with gut inflammation in PD, since inflammatory cytokines and activated glia cells were detected in both brain and intestine of PD patients (Cote et al., 2015; Devos et al., 2013; Morais et al., 2018). Sampson et.al found that gut microbes promoted a-synuclein-mediated motor dysfunction as well as reduced microglia activation(T. R. Sampson et al., 2016). Gut microbial imbalance leads to production of microbial metabolites such as SCFAs and lipopolysaccharide (LPS), which might be directly or indirectly involved in neuroinflammation due to their effects on gut–brain signaling pathways including immune system(Dalile et al., 2019). SCFAs, bacterial fermentation production, were found capable of modulating microglia-activation in central nervous system(CNS)(Erny et al., 2015), and having an impact on enteric nervous system (ENS) as well(Ono et al., 2004). Moreover, gliosis results in reduction in brain derived neurotrophic factor (BDNF) and glial cell derived neurotrophic factor (GDNF), which are derived from microglia and astrocytes to support the neuronal growth and survivals (Erickson et al., 2001). In addition, several studies have indicated that butyrate was correlative to gut barrier functions and gut inflammation (Kelly et al., 2015; Ploger et al., 2012), suggesting a link between gut microbiome and gut epithetical barrier.

Intestine hyperpermeability, which may result in a leaky gut in PD, is another pivotal aspect in microbiota-gut-brain axis, as porous epithelial barrier provides a way for neurotropic pathogens such as bacterial endotoxin LPS to initiate pathological process in ENS. Moreover, intestinal permeability of PD patients was found functionally dysregulated, and colonic occludin production, the tight-junction protein controlling intestinal barrier permeability, was observed decreased (Gershanik, 2018; Salat-Foix et al., 2012). Thus, we examined the alternation of zonula occluden-1 (ZO-1) and occudin in the colon and a clearly protein downregulation was observed in PD mouse model.

Furthermore, gut microbiota alternation and intestinal permeability change contribute to the activation of TLRs, which recognize molecules of microbial origin, i.e. microbial-associated molecular patterns (MAMPs) such as LPS, a microbial product and a potent TLR4 activator. TLRs are found presenting in the immune system and involved in innate immune responses. Previous studies have demonstrated that TLRs are expressed not only in neurons of the CNS but also in ENS, which are implicated in neurodegeneration and intestinal inflammation(Barajon et al., 2009; Okun et al., 2009; van Noort et al., 2009). There are evidences showing that LPS expressed on microglia and astrocytes was involved in the loss of dopaminergic neurons, and TLR4 could detect LPS leading to releasing of proinflammatory factors via activating nuclear factor kappa-B (NF-*κ*B) pathways(Anitha et al., 2012b; Castano et al., 1998). In patients with PD, TLR4 can activate downstream myeloid differentiation 88(MyD88)-dependent signal cascade mediating cytokines IL-1β level in colon(Perez-Pardo et al., 2019). More specifically, TLR4 recruits MyD88 signal adaptor protein, signaling through transcriptional NF-*κ*B or mitogen-activated protein kinases (MAPKs) pathways, which results in production of inflammatory cytokines like TNF-α and IL-1β. Interestingly, TLRs and MyD88 are widely expressed in the microglia astrocytes, oligodendrocytes and neurons of CNS(Hanke et al., 2011; Schroeder et al., 2021), and also present throughout the intestine(Anitha et al., 2012a; Ortega-Cava et al., 2003), indicating that TLR4/MyD88/NF-*κ*B might play a crucial role in neurodegenerative disease based on their immune properties on microbiota-gut-brain signaling, which was evidenced in our results that MPTP triggered inflammatory in gut and brain by activating TLR4/MyD88/NF-*κ*B signal cascades.

In current study, we demonstrated the disorders of gut-brain axis by gut microbial imbalance may play a pivotal role in pathogenesis of PD, which displayed as motor deficit, dopaminergic neuronal loss, gliosis, intense inflammatory in gut and brain, microbial dysbiosis, increased SCFAs productions, and enhancement of colonic epithelial permeability, and TLR4/MyD88/NF-*κ*B signaling pathway was indicated to be the underlying molecular mechanism mediating gut inflammation and neuroinflammation. However, CA exerted neuroprotective effects on PD mice by modulating microbiota-gut-brain axis.

## Results

### CA alleviates motor disorders in PD mice

To investigate the potential neuroprotective effects of CA on motor function in MPTP-induced PD mice, we subjected mice to a pole test for the assessment of bradykinesia by measuring the total descent time, and a traction test for the assessment of the muscle strength and equilibrium by scoring the traction. Compared to Control mice, mice in MPTP group exhibited motor dysfunction with longer descent time in pole test (Fig.1C, p<0.001) and lower scores in traction test (Fig.1D, p<0.01). However, CA+MPTP treated mice displayed a better motor performance compared to the MPTP group, which showed as the less time taken climbing-down the pole (Fig.1C, p<0.001) and the stronger muscle and better equilibrium gripping the rope (Fig.1D, p<0.01). Intriguingly, there were no obvious behavioral change between mice in CA60 and Control group, similarly, mice in CA30+MPTP and CA60+MPTP group had no difference in alleviating motor dysfunction, these results indicated that CA improved MPTP-induced Parkinsonian motor disorder.

**Fig.1.**
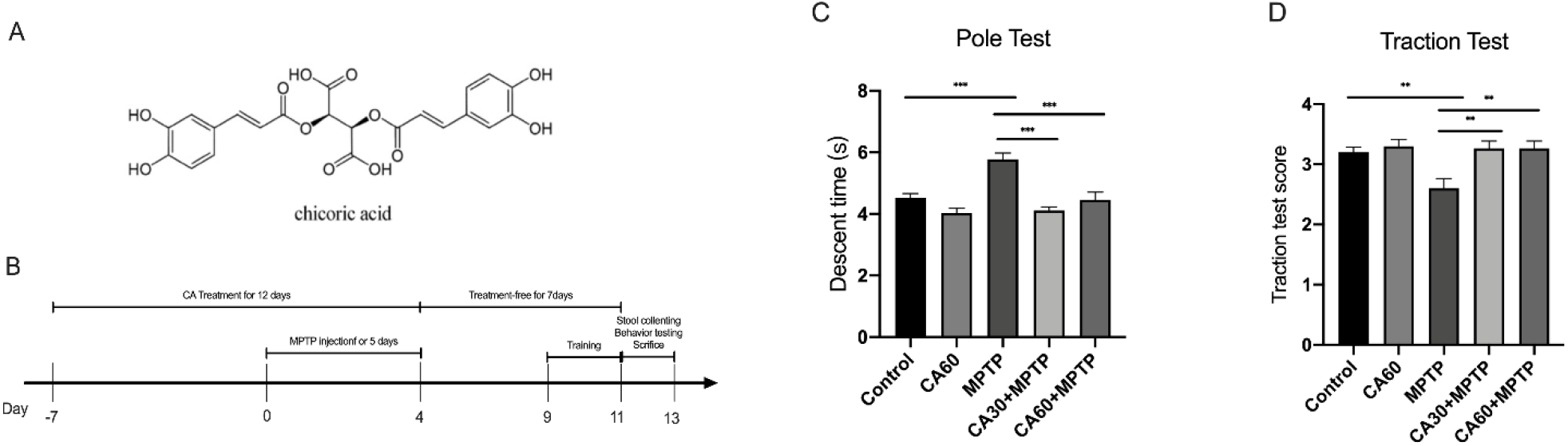
CA pretreatment improved the motor function in PD mice. A) Chemical structure of CA. B) Timeline for the experimental procedure: CA and MPTP Administration, behavior testing, stool sample and tissue collecting. C) Pole test: the pole descent time represents the degree of bradykinesia. Total descent time was significantly longer in MPTP mice compared with Control mice, and shorter in CA+MPTP mice compared with MPTP mice. D) Traction test: the suspension reflex score evaluated the muscle strength and equilibrium. The suspension reflex score was significantly lower in MPTP mice compared with Control mice, and higher in CA+MPTP mice compared with MPTP mice. Statistical comparison by one-way ANOVA with Tukey post hoc test; Value represent the means± SEM, **p<0.01, ***p<0.001. n = 12 mice per group.

### CA reversed the decline in dopaminergic neurons and striatal TH levels in PD Mice

To determine whether CA pretreatment prevents nigrostriatal DA lesion induced by MPTP. Survival of dopamine neurons in the SN was explored by immunofluorescence staining, and western blot analysis was applied to verify the immunofluorescence findings. As shown in Fig.2A and B, IF staining in the SN revealed a striking loss of TH-positive dopaminergic neurons in PD mice compared to Control mice (Fig.2A, p<0.001,), nevertheless, CA pretreatment dramatically rescued the MPTP-triggered loss of dopaminergic neurons in CA+MPTP group (Fig.2A, p<0.05). Western blotting of striatal tissue showed a clearly lower level of TH-expression in MPTP treated mice compared to Control mice, whereas, a marked enhancement of TH-expression was observed in CA+MPTP group (Fig.2C and D. p<0.05). In addition, Control mice exhibited the nearly equal numbers of dopaminergic neuron and expression of TH to CA60 mice. Likewise, there were no significant differences between CA30+MPTP and CA60+MPTP group in alleviating dopaminergic neuron loss in SN and TH-expression decline in striatum. These results suggested that prophylactic administration of CA can effectively prevent MPTP-induced loss of dopaminergic neurons in the SN and the decrease of striatal TH level.

**Fig.2.**
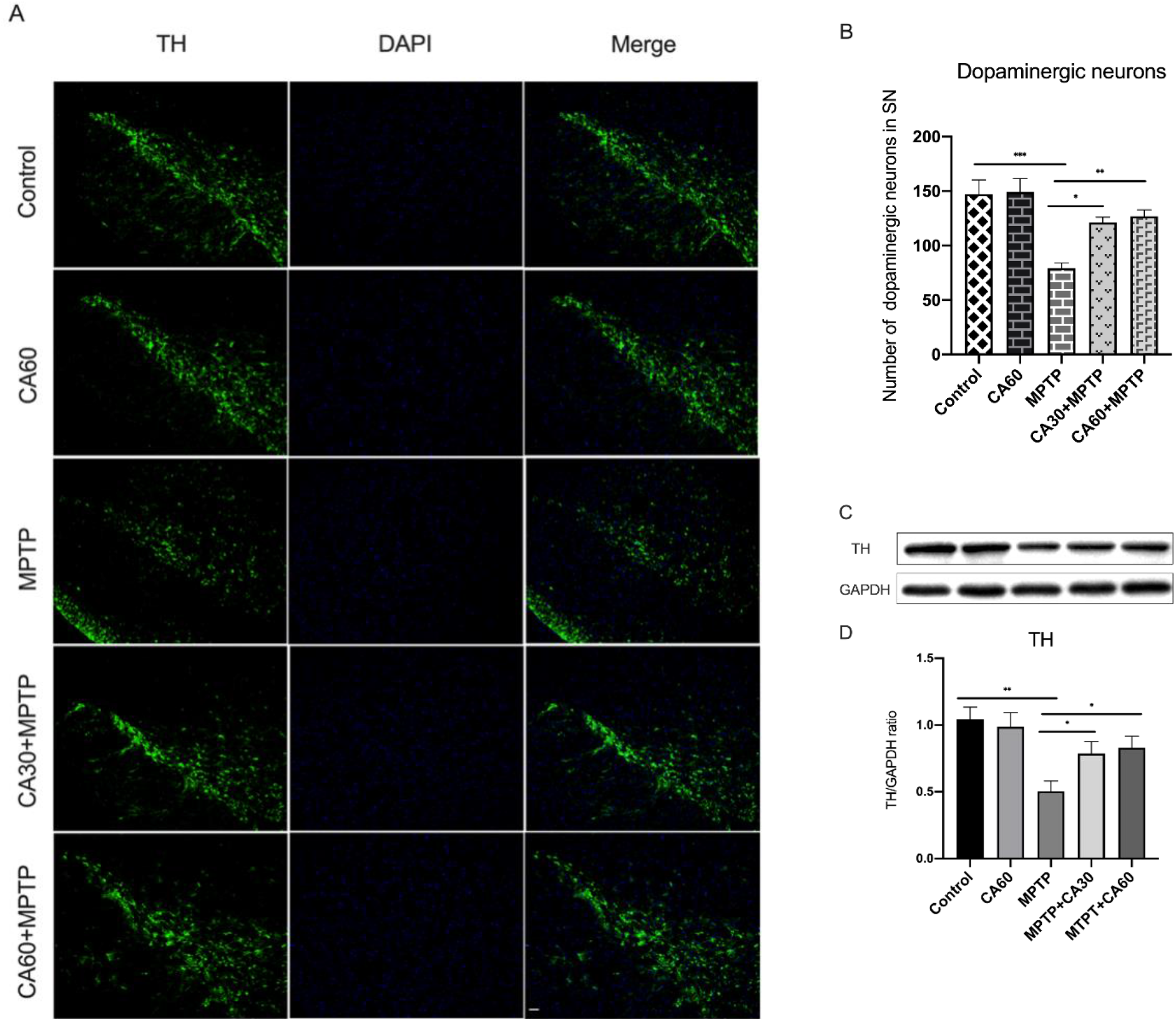
CA pretreatment promoted the dopaminergic neurons survival in the SN and TH expression in striatum. (A) Representative immunofluorescence staining for TH in the SN. Scale bar is 100µm. B) Quantitative analysis for the number of TH immunoreactive cells. (C) Representative bands of western-blot for TH in striatum. (D) Relative protein level of striatal TH. Quantification: band intensity normalized to GAPDH. Statistical comparison by one-way ANOVA with Tukey post hoc test; values are represented as the means ± SEM. *p < 0.05, **p < 0.01, ***p<0.001. n = 6 mice per group.

### CA attenuated MPTP-indued glial activation accompanied by restoration of BDNF and GDNF

Glial cells and neurotrophic factors, involved in neuroinflammation and neurodegeneration of dopaminergic neurons, are essential for maintaining nervous system function(Palasz et al., 2017; Xu et al., 2017). we conducted immunofluorescence staining against GFAP and Ibal-1, the markers of activated astrocytes and microglia, respectively. As shown in Fig.3A and Fig.3C, alone with the reduction of TH-positive cells, the population of astrocytes and microglia in the SN were sharply activated after MPTP challenge, as evidenced by increased production of GFAP and Ibal-1(Fig.3B and 3D, p<0.001). However, this gliosis indued by MPTP was attenuated with CA pretreatment, as evidenced by the significantly decreasing production of GFAP (Fig.3B, p<0.01) and Ibal-1(Fig.3B, p<0.05 and P<0.01,) in the CA+MPTP group, suggesting CA maintained glia population in PD mice close to Control mice. To further ascertain CA has an effect on neurotrophins in PD mice following gliosis, we also detected levels of BDNF and GDNF in striatal tissue by western blot analysis (Fig.3E and 3G). The results indicated the depletion of BDNF (Fig.3F, p<0.001) and GDNF (Fig.3H, p<0.01) in MPTP-induced mice compared to Control mice, whereas, mice in CA +MPTP group exhibited a distinctly restoration of BDNF (Fig.3F, p<0.01) and GDNF (p<0.01, Fig.3H) compared to MPTP group, suggesting this neurotrophins-suppression caused by MPTP was effectively prevented by CA pretreatment. Additionally, CA60 mice were indistinguishable from Control mice in regard to population of glia population and neurotrophins (BDNF and GDNF) levels, similarly, these changes were not significant between CA30+MPTP and CA60+MPTP groups. The results above revealed that CA had a marked inhibition on the increasement of brain glia population and the depletion of BDNF and GDNF in PD mice.

**Fig.3.**
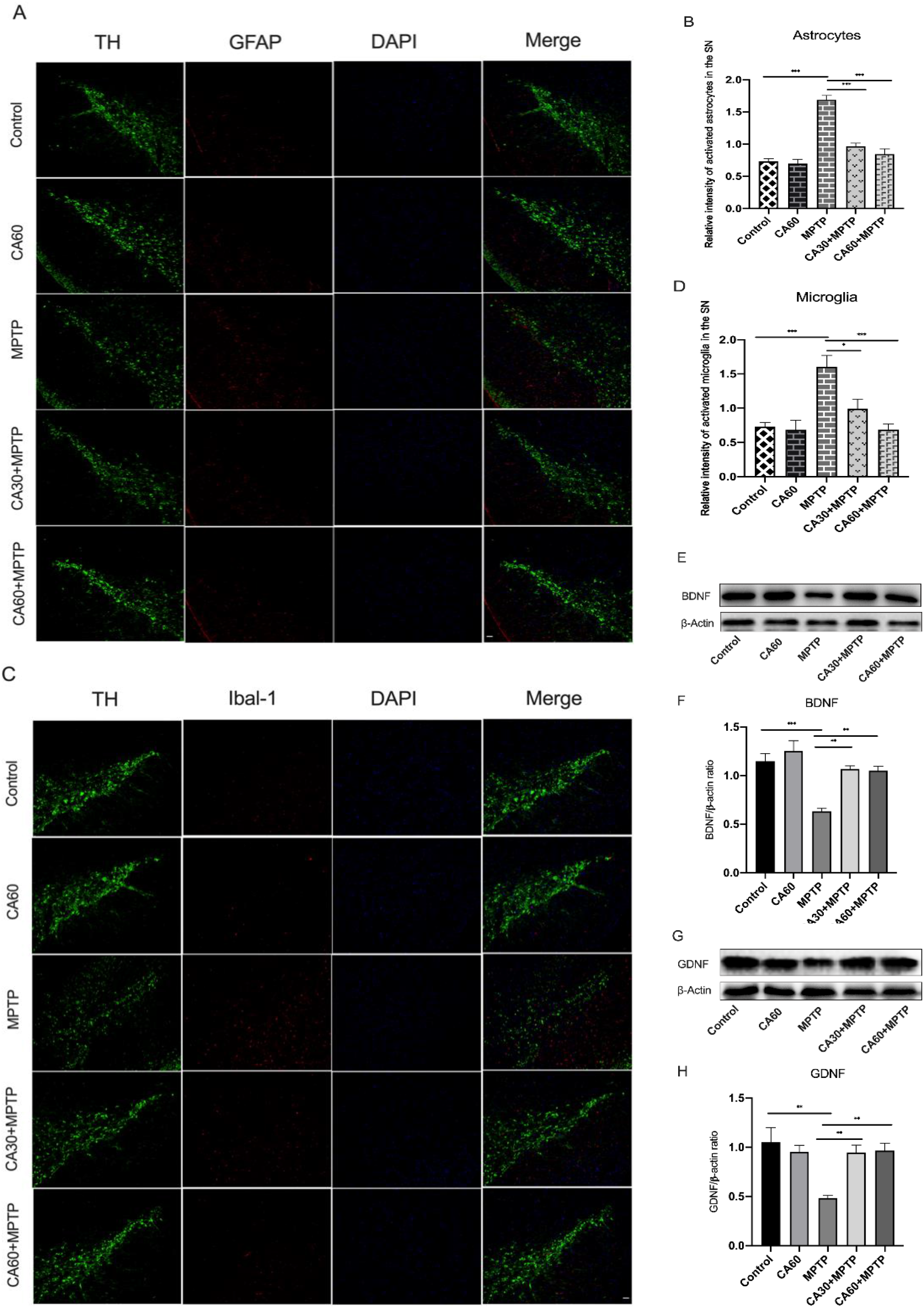
CA attenuated MPTP-indued glial activation accompanied by restoration of BDNF and GDNF. Scale bar is 100µm. (A) Representative double-immunofluorescence staining for TH (green) and GFAP (red) in the SN. Scale bar is 100µm. (B) Quantitative analysis for relative density of GFAP positive cells. (C) Representative double-immunofluorescence staining for TH (green) and Ibal-1(red) in the SN. (D) Quantitative analysis for relative density of Ibal-1 positive cells. (E,G) Representative bands of western-blot for Striatal BDNF and GDNF, respectively. (F, H) Relative protein level of Striatal BDNF and GDNF, respectively. Quantification: band intensity normalized to β-Actin. Statistical comparison by one-way ANOVA with Tukey post hoc test; values are represented as the means ± SEM. *p < 0.05, **p < 0.01, ***p<0.001. n = 4 mice per group.

### CA exerted an influence on SCFAs concentration, and ameliorated colonic epithelial barrier impairment in PD mice

Increasing evidences in favor of SCFAs being the major mediators regulating the microbiota-gut-brain axis. Therefore, we performed Fecal SCFAs concentrations detection via GC–MS analysis. We detected various SCFAs, of which six SCFAs were detected. As exhibited in Fig.4, acetic, propionic, isobutyric, butyric, isovaleric, valeric acid were overall increased in MPTP mice compared to Control mice(Fig.4A,p<0.001;Fig.4B,p<0.05; Fig.4C,p<0.001;Fig.4D,p<0.001;Fig.4E,p<0.001;Fig.4F,p<0.001), however, CA pretreatment effectively restored the level of SCFAs in MPTP-treated mice, excluding propionic acid (Fig.4A,p<0.05 and p<0.01;Fig.4C,p<0.05 and p<0.01;Fig.4D,p<0.001 and p<0.001;Fig.4E,p<0.001 and p<0.001;Fig.4F,p<0.01 and p<0,01), but the concentration-change of these SCFAs between CA30+MPTP group and CA60+MPTP group was not significant. Furthermore, mice in CA60 group exhibited a higher degree of butyric acid compared to Control mice (Fig.4D, p<0.001). These data suggested that CA had an influence on fecal SCFAs content in PD mice of which gut microbiota was altered. In addition, we examined the histological changes of colonic epithelial barrier and levels of colonic zonula occludens-1 (ZO-1) and occludin. Compared to Control mice, PD mice displayed the loss of crypts and glands, edema of the lamina propria, and thinning of the muscularis mucosa, epithelium and intestinal mucosa (Fig.5A), moreover, the lower expression of ZO-1(Fig.5C, p<0.001) and occludin (Fig.5D,6<0.001) was observed in MPTP group. However, CA pretreatment prevented the substructural damage to the colonic mucosa, similarly, the decrease in protein levels of ZO-1(Fig.5C, p<0.01 and p<0.01) and occluding (Fig.5D, p<0.01 and p<0.05) was clearly reversed in CA+MPTP groups, Furthermore, there were no significant differences in changing colonic mucosa as well as expression of ZO-1 and occludin between Control and MPTP groups, consistently, these changes were not significant between CA30+MPTP and CA60+MPTP groups. In summary, CA favorably alleviated the abnormal SCFA-increasement in PD mice and ameliorate colonic epithelial barrier impairment caused by MPTP.

**Fig.4.**
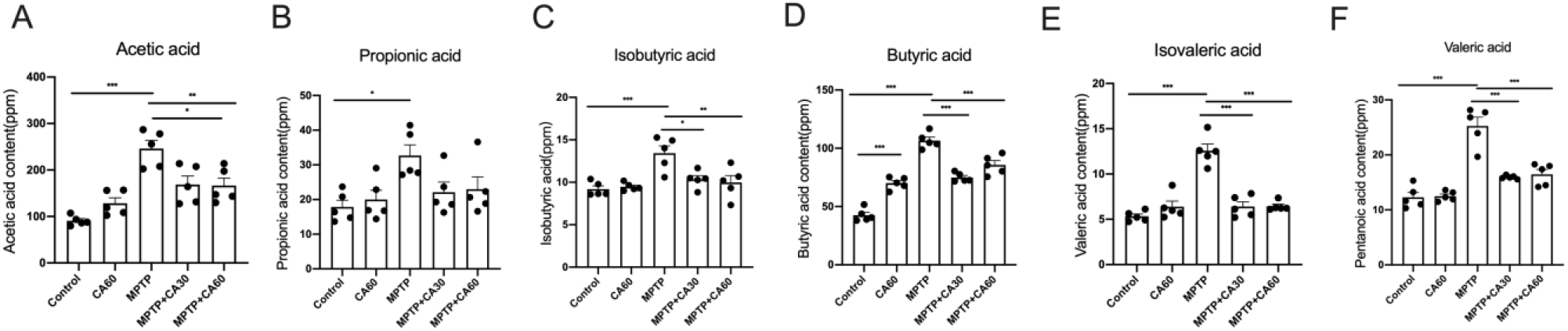
Fecal SCFAs alterations in mice. (A) Fecal acetic acid content. (B) Fecal propionic acid content. (C) Fecal isobutyric acid content. (D) Fecal butyric acid content. (E) Fecal isovaleric acid content. (F) Fecal valeric acid content. Statistical comparison by one-way ANOVA with Tukey post hoc test; values are represented as the means ± SEM. *p < 0.05, **p < 0.01, ***p<0.001. n = 5 mice per group.

**Fig.5.**
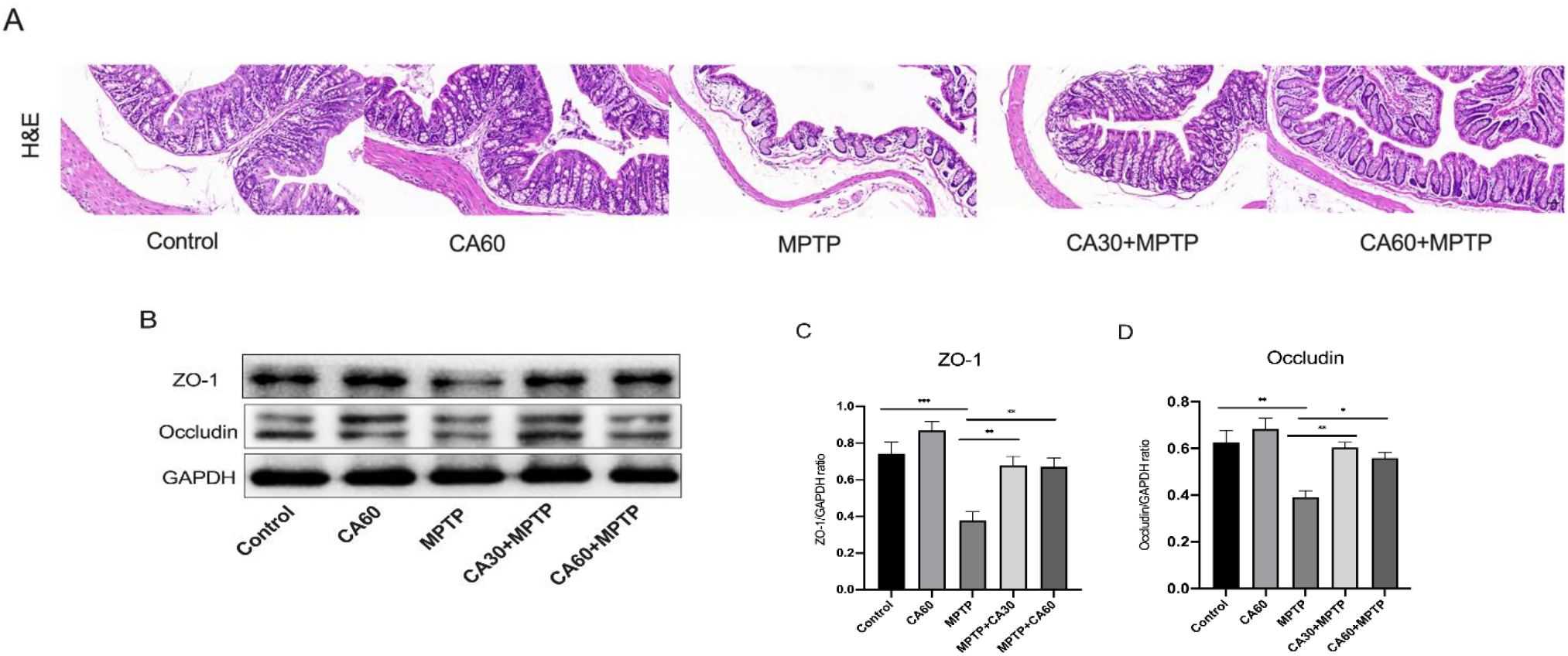
CA ameliorated colonic epithelial barrier impairment. (A) Colon histopathology. Scale bar is 50μm. (B)Representative bands of western-blot for colonic ZO-1 and occludin, respectively (D) Relative protein level of colonic ZO-1 and Occludin, respectively. Quantification: band intensity normalized to GAPDH. Statistical comparison by one-way ANOVA with Tukey post hoc test; values are represented as the means ± SEM. *p < 0.05, **p < 0.01, ***p<0.001. n = 4 mice per group.

### CA modified the gut microbial dysbiosis induced by MPTP

We assessed the gut microbiota using 16s rRNA sequencing and bio-informatic analysis, which focus on abundance, diversity and distribution of gut microbiota and demonstrated a holistic status of gut microbiota. Based on alpha diversity analysis which is indicated by Chao1 index and used to estimate the richness of microbial community richness, as displayed in Fig.6, a significant increasement of Chao1 index was observed in the MPTP group compared to Control group, and no difference between MPTP group and CA30+MPTP group, however, Chao1 index was observably decreased in CA60+MPTP group which was approximated to the Control group. Additionally, there was no significant difference in α-diversity between Control group and CA60 group (Fig.6A). Beta diversity was further used to evaluate the extent of the similarity of gut microbial communities between groups using PCoA based on the weighted UniFrac distance at the OTU level. As shown in Fig.6B, the overall structures of the bacterial communities in PD mice were significantly different from the mice in Control, CA60, CA30+MPTP and CA60+MPTP group, but there were no significant differences between 4 groups. However, compared to CA60+MPTP group, bacterial compositions in CA30+MPTP were closer to the Control group, suggesting that CA treatment could alter the microbiota compositions approximately to normal level (Fig.6B).

**Fig.6.**
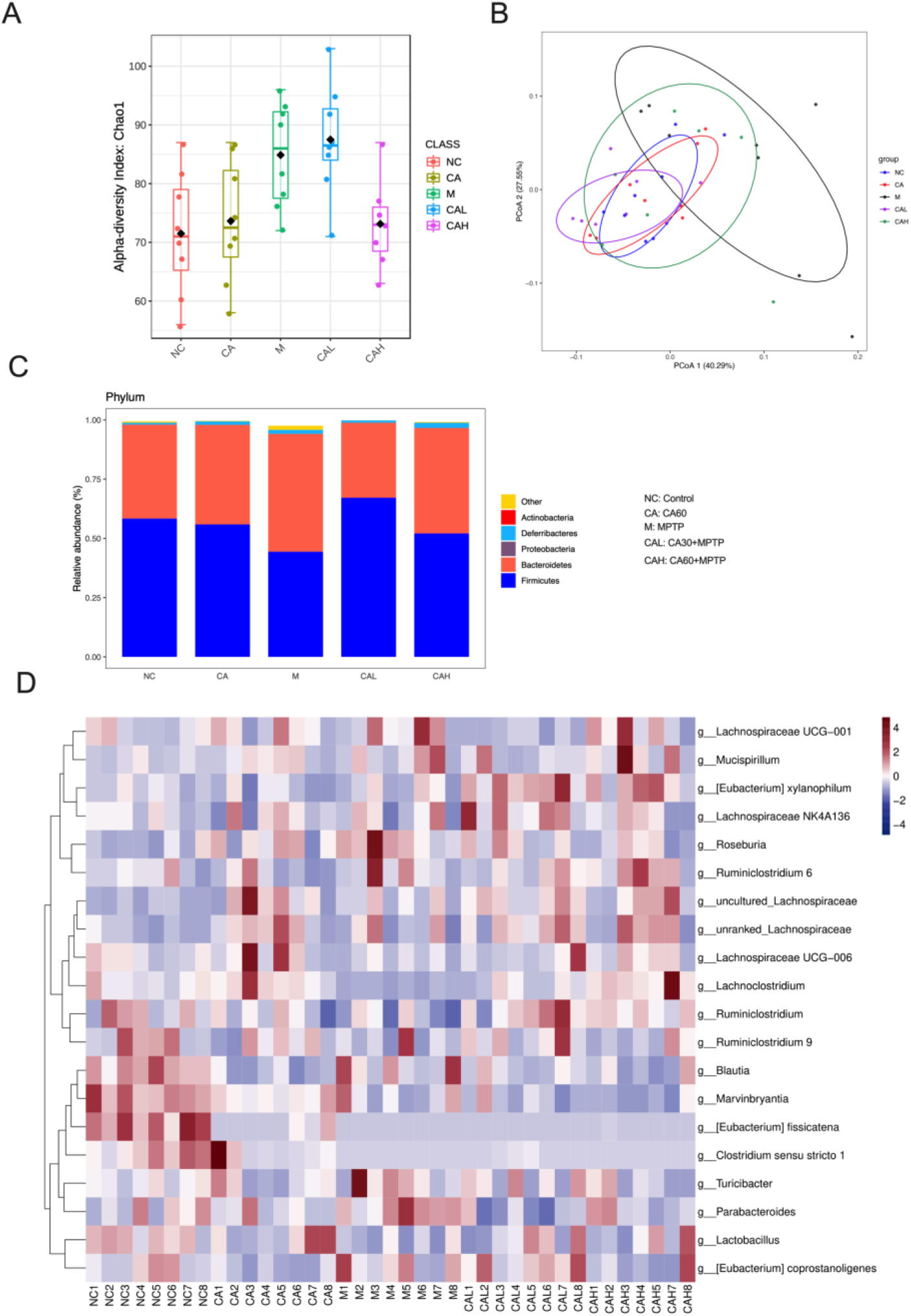
CA and MPTP treatment affected the mouse fecal microbiomes based on analysis of 16S rRNA sequencing. (A) Alpha diversities of bacteria communities in individual samples from each group shown by Chao-1 index. (B) Principal coordinate analysis (PCoA) plots based on weighted UniFrac metrics of bacterial diversity. (C) The average relative abundance of the microbial community for each group at the phylum level. Only those bacterial phyla that comprised on average >1% of the relative abundance across all samples are displayed. (D) The heatmap of the relative abundance of top 20 gut microbiota at genu level. C=Control group, CA=CA60 group, M=MPTP group, CAL=CA30+MPTP, CAH=CA60+MPTP. n=8 mice in Control group, CA60 group, MPTP group and CA30+MPTP group, and n = 7 mice in CA60+MPTP group

Furthermore, a number of altered gut microbiota were identified at the phylum level, only those bacterial phyla averaged >1% of the relative abundance across all samples were displayed, the bar-plots revealed a clear alternation of relative abundance of phylum-level microbiota in MPTP group, shown as a higher abundance of *Bactroidetes* and a lower abundance of *Fimicutes*, which in CA+MPTP groups tended to be decreased and increased, respectively, moreover, the ratio of phylum *Fimicutes/Bactroidetes* of CA30+MPTP were higher than that of CA60+MPTP, and the latter was approximated to that of Control group (Fig.6C). In addition, a total of 60 genera were identified across the five groups and the relative abundance of top 20 gut microbiota at genu-level were displayed in *heatmap* (Fig.6D). Genus-level analyses indicated the distinction of gut microbiota between the five groups. More specifically, as shown in *heatmap*, compared to Control and CA treated groups, an increasing trend of genu relative abundance for *Parabacteroide, Roseburia* and *Turicibacter* and a reduced trend of genu relative abundance for Lactobacillus, *Riminiclostridium, Lachnoclostridium* were found in MPTP group, however, these changes were not significant, suggesting the sample size is required to be expanded in further study. Overall, results here revealed a presence of gut microbial imbalance in MPTP-induced mice, and CA treatment appeared a favor modification of gut microbial dysbiosis.

### CA inhibited inflammation both in gut and brain possibly by suppressing TLR4/MyD88/NF-*κ*B signaling cascade

To further explore the interaction mechanism of CA on neuroinflammation and gut microbial dysbiosis in MPTP-triggered PD, we characterized the expressions of TLR4, MyD88 and NF-*κ*B in striatum and colon by Western-blot analysis (Fig.7A and Fig.7E), and TNF-α and IL-1β in serum, striatum and colon by ELISA (Fig.7I∼N). Notably, compared to Control group, the levels of TLR4(Fig.7B,p<0.01 and Fig.7F,p<0.001), MyD88 (Fig.7C,p<0.01 and Fig.7G,p<0,001) and NF-*κ*B(Fig7D,p<0.01 and Fig.7H,p<0.001) in the striatum and colon were enhanced dramatically after MPTP challenge, a marked upregulation of TNF-α was also observed(Fig.7J,p<0.01,Fig.7K,p<0.05 and Fig.7M,p<0.001)and IL-1β (Fig.7K,p<0.01,Fig.7L,p<0.01 and Fig.7N,p<0.001)in serum, striatum and colon of MPTP group, nevertheless, these enhancements of protein expression and inflammatory cytokine content were significantly restrained with CA pretreatment in PD mice, which showed as downregulated levels of TLR4(Fig.7B,p<0.05 and p<0.01;Fig.7F,p<0.01 and p<0.01), MyD88 (Fig.7C,p<0.05 and p<0.01;Fig.7G,p<0.05 and p<0.001) and NF-*κ*B(Fig.7D,p<0.01 and p<0.01; Fig.7H,p<0.05 and p<0.05), as well as decreased concentration of TNF-α (Fig.7I,p<0.05 and p<0.01;Fig.7K,p<0.05 and p<0.01; Fig.7M,p<0.01 and p<0.05) and IL-1β(Fig.7J,p<0.01 and p<0.01; Fig.7L,p<0.01 and p<0.05, Fig.7N,p<0.001 and p<0.001) in CA+MPTP groups, moreover, compared to CA30+MPTP group, mice in CA60+MPTP group displayed a stronger inhibition on activation of striatal TLR4 and colonic TLR4 and MyD88, but indistinctive difference in level of TNF-α and IL-1β in colon and striatum. Intriguingly, there were no obvious changes in expression of TLR4, MyD88 and NF-*κ*B and concentration of TNF-α And IL-1β in the striatum and colon between Control and CA60 group. These findings highlight the fact that the TLR4/MyD88/ NF-*κ*B signaling pathway played a key role in PD mice accompanied with neuroinflammation and gut inflammation, and CA pretreatment may improve the gut microbiome-dysbiosis to inhibit neuroinflammation and gut inflammation by suppressing this signaling cascade.

**Fig.7.**
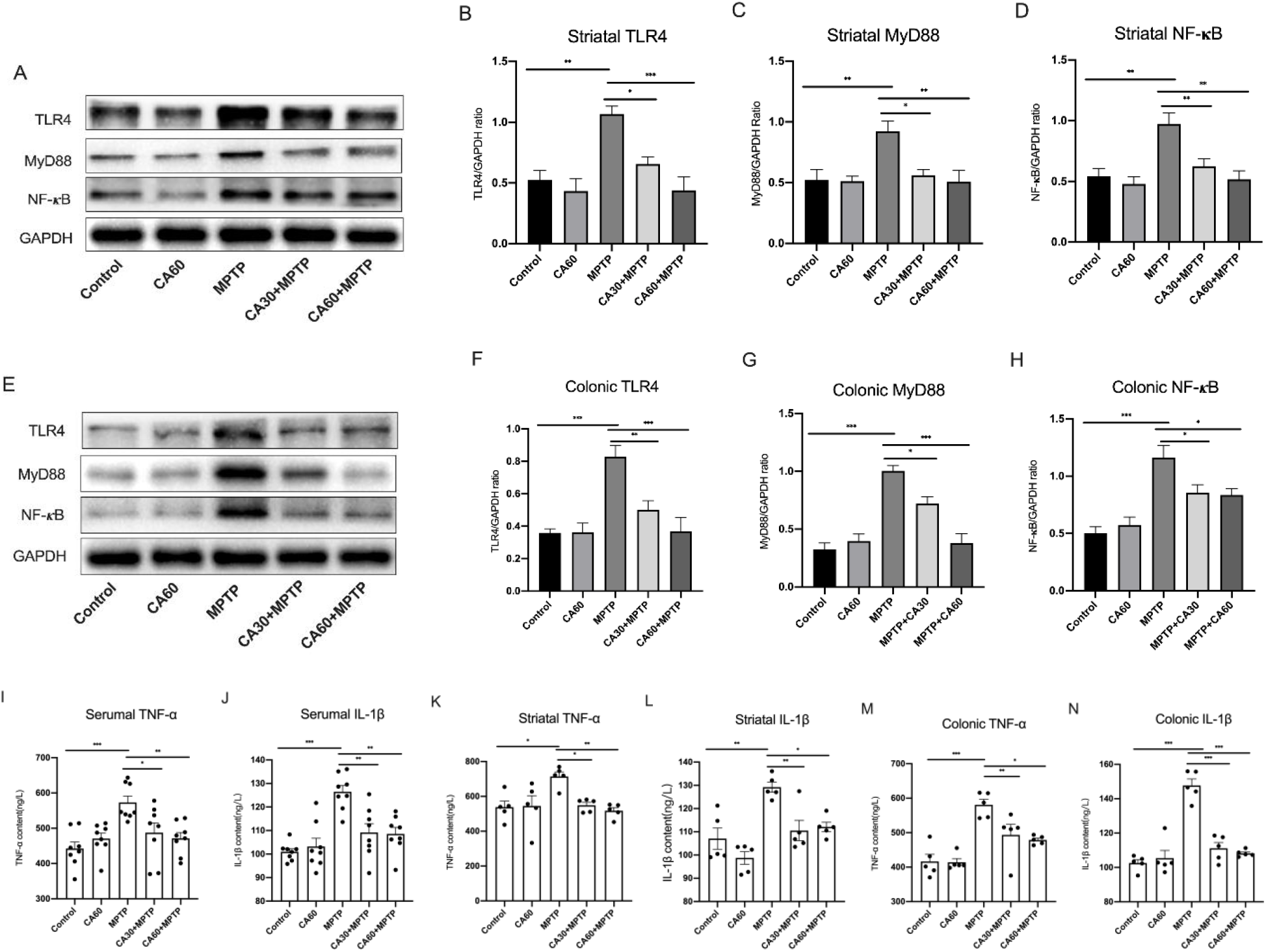
(A) Representative bands of western-blot for striatal TLR4, MyD88 and NF-*κ*B, respectively. (B-D) Relative protein level of striatal TLR4, MyD88 and NF-*κ*B, respectively. Quantification: band intensity normalized to GAPDH. (E) Representative bands of western-blot for colonic TLR4, MyD88 and NF-*κ*B, respectively. (F-H) Relative protein level of colonic TLR4, MyD88 and NF-*κ*B, respectively. Quantification: band intensity normalized to GAPDH. (I, K and M) The concentration of TNF-α in the serum, striatum and colonic, respectively. (J, L and N) The concentration of IL-1β in the serum, striatum and colonic, respectively. Statistical comparison by one-way ANOVA with Tukey post hoc test; values are represented as the means ± SEM. *p < 0.05, **p < 0.01, ***p<0.001. n = 4 mice per group for western blot analysis, n=8 mice per group for serum ELISA, n=5 mice per group for striatum and colon ELISA.

## Discussion

The imbalance of the gut-microbiome-brain axis by gut microbial dysbiosis is a crucial factor in pathogenesis and development of PD (Baizabal-Carvallo et al., 2020). Chao1 index represents the alpha diversity of the microbial community, higher Chao1 index signifies an increase in microbial community. In this work, the alpha diversity of gut microbiota was increased in the MPTP mice with a higher Chao1 index, nevertheless, CA pretreatment at high dosage deterred MPTP-induced ascent of alpha diversity. On the basis of PCoA, the bacterial compositions of mice pretreated by CA approximated to that of Control mice. Interestingly. Although the alpha diversity of CA30+MPTP group pretreated by CA at low dosage changed insignificantly, the bacterial compositions in CA30+MPTP group were closer to the Control group compared to CA60+MPTP group. Combing with the impacts of CA-pretreatment at low dosage on gut-brain axis partially, we inferred that CA-pretreatment at low dosage had no effect on alpha diversity of PD mice, but well restored the structural changes of the bacterial communities caused by MPTP, and strikingly increased the ratio of phylum *Fimicutes*/*Bactroidetes*, suggesting that gut bacterial compositions were heavily involved in anti-inflammation and dopaminergic neuronal survivals. Together, the present results revealed that CA is capable to prevent the MPTP-induced increasement in alpha diversity and dissimilarity in beta diversity. Further analysis on microbial compositions here demonstrated that PD mice possessed a lower abundance of *Firmicutes* as well as higher abundance of *Bacteroidetes* at phylum level. A similar phenomenon was also confirmed in Sun’s work that a phylum Firmicutes was decreased but *Bacteroidetes* changed indistinctively in murine model of PD triggered by MPTP (M.-F. Sun et al., 2018). Of note, gut microbial dysbiosis with decreased *Firmicutes* and increased *Bacteroidetes* was also observed in Alzheimer’s disease (AD) patients (Vogt et al., 2017). Moreover, a high level of *Bacteroidetes* is found in autistic individuals (Finegold et al., 2010), suggesting that the phyla *Firmicutes* and *Bacteroidetes* were involved in nervous system disorder via gut-microbiome-brain axis. Genus-level comparison analysis on the specific taxa was performed to assess the global microbiota changes. Intriguingly, here escalating trend of *Parabacteroide* belonging to phylum *Bacteroidetes* and the decline trend of phylum *Firmicutes* were observed following decreased Lactobacillus, *Riminiclostridium* and *Lachnoclostridium* but conversely increased *Roseburia* and *Turicibacter*, in order to confirm the genus-level changes of gut microbiota, the sample size should be expanded. Taken together, it was speculated that CA has the potential for anti-inflammation and neuroprotection by deterring MPTP mediated changes in gut microbiome. Nevertheless, further studies are required to elucidate how the alternation of microbiome in the gut is associated with the neuroprotection of CA in the brain.

SCFAs, the most detected microbial-derived metabolites, of which more than 95% consists of acetate, propionate, and butyrate, can not only serve as a gut microbial mediator to regulate the intestinal immune function including inflammation, but also transport across the BBB to exert an influence on microglia (Braniste et al., 2014; Cryan et al., 2019; Timothy R. Sampson et al., 2016; Vinolo et al., 2011). Sampson et al. observed the exacerbated α-synuclein-related neurodegeneration and neuroinflammation in PD mice being received the cocktail of acetate, propionate and butyrate by oral administration(Timothy R. Sampson et al., 2016), indicating an interplay between SCFAs and neuroinflammation in the brain. In this study, the detected six SCFAs of which five were overall increased in MPTP mice, leading to a suggestion that increasement of SCFAs caused by gut microbial imbalance might have the positive correlation with the activated microglia and astrocytes, as SCFAs were decreased while gliosis was inhibited in current observations. In addition, the results that higher degree of butyric acid in MPTP mice might result from the enhanced abundance of *Roseburia*, which is the producer of butyrate (Westfall et al., 2017). The striking increase of valeric acid concentration in MPTP mice was also found in previous reports (M. F. Sun et al., 2018; Zhou et al., 2019).

Gut microbial imbalance was heavily involved in colonic mucus barrier(Jakobsson et al., 2015). Bacterial products such as LPS could induce increases of permeability of gut epithelial barrier and BBB, and release of proinflammatory factors(Kesika et al., 2021; Villaran et al., 2010). Additionally, alternation of SCFAs might have the interactions with gut epithelial barrier function(Topping et al., 2001). Recent reports suggested that MPTP could cause gut epithelial barrier impairment with a disorganization of proteins ZO-1 and occludin in the colon(Huh et al., 2020). Since we observed MPTP-triggered microbial imbalance in gut and abnormal generation in SCFAs, together with higher degree of TNF-α and IL-1β in the colon, brain and serum of PD mice, it was inferred that MPTP might mediate gut inflammation and neuroinflammation accompanied with colonic epithelial barrier impairment, which was evidenced by as amelioration of colonic epithelial barrier disruption and significant down-regulation of ZO-1 and occludin in the colon of PD mouse in this study. Most notably, CA pretreatment restored the abnormal elevation of SCFAs and dramatic reduction in colonic tight junction proteins. Considering that CA modulated the gut microbiota compositions and reduce the TNF-α and IL-1β in the colon, brain and serum of PD mice, it is reasonable to speculate that CA might regulate SCFAs production and colonic permeability to alleviated gut inflammation and neuroinflammation by shaping gut microbiome. Likewise, how CA specifically works in gut-brain axis of PD need to be further explored.

As is well-known, PD is positively correlative to a massive microgliosis and astrogliosis, which trigger and amplify neuroinflammation by releasing inflammatory cytokines such as TNF-α and IL-1β. Furthermore, activated glial cells possess neurotrophic properties, in which both BDNF and GDNF are acknowledged to promote the growth and survival of dopaminergic neurons(Imamura et al., 2005). Also, a growing number of studies revealed the interplay between neurotrophic factors and glial population. Rajkumar Verma et al. found that the decline in BDNF expression showed possible correlation to the higher degree of TNF-α and IL-1β(Verma et al., 2017). Consistently, TNF-α displayed an inhibition on GDNF levels in a recent report (Di Persio et al., 2020), and IL-1β compromised neuronal survival by blocking the production of BDNF (Tong et al., 2008). In the present study, CA pretreatment downregulated the productions of GFAP and Ibal-1 in the SN, correspondingly, CA prevented the reduction in striatal BDNF and GDNF coupled with a clearly decreased concentration of TNF-α and IL-1β in the striatum and serum. Combined with previous studies and current findings, we came up with an inference that CA can restrain the activations of gliosis and enhanced the levels of BDNF and GDNF to protect dopaminergic neurons against MPTP through inhibiting the generation of TNF-α and IL-1β. Notably, the inflammatory cytokines might be simultaneously released from leaky gut and activated glial population, where the serious inflammatory response was observed.

Compelling evidences suggest a link between gut microbial, neuroinflammation and TLR4. Previous study in PD mouse model displayed a reduction in dopamine and an alleviation of neuroinflammation under the absence of TLR4(Campolo et al., 2019). Anitha et.al revealed that lipopolysaccharide, a gut microbial product, appeared to have an impact on neuronal survival by modulating inflammation via TLR4 signaling.(Anitha et al., 2012b). In another report, the inflammatory polysaccharide from gut bacterium *Ruminococcus gnavus*, induced the secretion of inflammatory cytokines like TNF-α, which aggravated the intestinal inflammatory response (Henke et al., 2019). In addition, the gut microbiota was able to induce macrophages and dendritic cells to produce IL-1β, another amplifier of inflammation(Rosser et al., 2014; Shaw et al., 2012). Above reports suggested that TLR4 pathway might play a pivotal role in PD pathogenesis by driving gut microbiota-mediated gut inflammation and neuroinflammation. In present results, TLR4/MyD88/NF-kB in PD mice appeared a higher level in colon and striatum, whereas CA pretreatment reduced the signaling molecules expression to alleviated the inflammation in gut and brain. Combining with other results in current study, we speculated that CA might minimize neuroinflammation and support neuronal survival by modulating the microbial dysbiosis-driven inflammation in gut via TLR4/MyD88/NF-*κ*B signaling pathway. Also, there might exist other signaling pathways involved in inflammation reactions of PD mice, as MyD88 is a pivotal adaptor not only in TLRs signaling pathway but also IL-1R signaling pathway, NF-*κ*B, the downstream effector of both TLRs/MyD88 and IL-1R/MyD88 pathways, contributes towards the increases of inflammatory cytokines and chemokines while being activated(Hanke et al., 2011; Xiong et al., 2020). Multiple mechanisms and pathways by which the interplay between brain and gut influence inflammatory diseases of the central nervous system where signaling molecules like TLR4 and MyD88 are widely expressed (Grenham et al., 2011; Hanke et al., 2011). In recent years, evidences have shown that CA via TLR4/NF-*κ*B pathway plays the anti-inflammatory activity including neuroinflammation. Nonetheless, limited studies focused on gut-microbiome-brain axis, through which CA is capable to inhibit neuroinflammation and protect dopaminergic neurons against MPTP.

In summary, the present findings demonstrated that oral administration of CA provided a neuroprotection or neurorescue against MPTP-induced DA neuronal impairment via gut-microbiome-brain axis. The possible mechanism found in current study showed that CA ameliorate the inflammation in both brain and gut throughout downregulating the TLR4/MyD88/NF-*κ*B signaling cascade.

## Materials and Methods

### Animals and treatment

All male C57BL/6 mice used in this study were acquired from Beijing Weitonglihua Laboratory Animal Technology Institute (Shanghai, China). Animals were adapted to feed for a week before the formal experiments and kept in a standard feeding environment (pathogen-free, 24±2°C, 55 ± 10% humidity, 12-12hr light/dark cycle) with food and water ad libitum. Animal welfare and experimental procedures were conducted in accordance with the NIH Guide for the Care and Use of Laboratory Animals (NIH, revised 1996) and approved by the Animal Ethics Committee of Jiangnan University (Jiangsu, China). Sixty eight-week-old mice (male,18g±2g) were randomly divided into five groups (12 mice per group). The control group was treated with normal saline, CA60 group was treated with CA (Shanghai yiyan Bio-technology, China) by oral gavage at dose of 60mg/Kg/d, MPTP group was treated with MPTP (Sigma-Aldrich) by intraperitoneal injection at dose of 30mg/Kg/d, CA30+MPTP and CA60+MPTP groups were treated with CA by oral gavage at dose of 30mg/Kg/d and 60mg/Kg/d, respectively, then MPTP by intraperitoneal injection at dose of 30mg/Kg/d.

Starting on the first day of the experiment, mice in the CA60, CA30+MPTP and CA60+MPTP were received CA (dissolved in saline) for 12 consecutive days, the other two groups received normal saline in the same volume. From day8 to day12 of the experiment, 2hr after CA administration, mice in the MPTP group, CA30+MPTP and CA60+MPTP were received MPTP (dissolve in saline) injection for 5 consecutive days, the other two groups were received normal saline in the same volume. After the last MPTP injection, no further treatments were given to any of the mice for 7 subsequent days. Feces collection and behavioral experiments were carried out in next 2 days. Subsequently, the mice were humanely killed after anesthetized with isoflurane, serum, striatum and colon tissue were collected and immediately stored at −80°C for further usage, the whole brain was removed for frozen section after the mice were transcardially perfused with 0.01M of phosphate buffered saline (PBS) followed by 4% paraformaldehyde (PFA) (w/v) in 0.01M of PBS (pH=7.4). This experimental timeline is illustrated in Fig.1B.

### Behavioral test

The behavioral test was carried out as previously described(Kuribara et al., 1977; Luchtman et al., 2009). On the 5th day of 7days-treatment-free, the mice were received daily behavioral training for 3 days. The formal behavioral testing was performed the next day.

### Pole test

A 1 cm in diameter and 50 cm in height wooden pole, with a spherical protuberance, 2 cm in diameter on top, was entangled with non-adhesive gauze and vertically fixed in the home cage. The mouse was placed head-down on the top of the pole, the total time for each mouse to reach the base of the home cage was recorded. The test for each animal was repeated three times at 10-min intervals, the average time was taken for statistical analyses.

### Traction test

The fore limbs of the mice were placed on a straight and horizontal rope (diameter 5 mm) while the hind limbs were placed without restriction, the condition of gripping the rope with limbs were observed for 10s and scored from 1 to 4: 1 point for gripping the rope with one front paws, 2 points for gripping the rope with both front paws, 3 points for gripping the rope with one hind paw, 4 points for gripping the rope with both hind paws. The test for each animal was performed three times and the average score was taken for statistical analyses.

### Enzyme-Linked immunosorbent assay (ELISA)

The content of TNF-α and IL-1β in serum, striatum and colon was detected by commercial ELISA kit (Mouse TNF-α Kit, Mouse IL-1β Kit) (Nanjing Jiancheng Bioengineering Institute, China). All experimental procedures were conducted according to the manufacturers’ protocol. TNF-α and IL-1β concentration were expressed in ng/L protein.

### Immunofluorescent staining and image analysis

The whole brain was removed and post-fixed in 4% PFA at 4°C for 24 h, then dehydrated in 20% sucrose solution at 4°C until the brain sank to the bottom of the container, then 30% sucrose solution until subsidence at 4°C, and then embedded in optimal cutting temperature compound (SAKURA, USA) for frozen sectioning containing substantia nigra pars compacta (SNpc), each mouse brain was cryosectioned at 10μm by cryostat microtome (Leica, CM1950, Germany).

Brain slices were immersed in 0.01M sodium citrate buffer (pH 6.0) for antigen retrieval and washed three times in PBS for 10min each time. Then, the slices were incubated in PBS containing 0.2% Triton X-100(v/v) and 5% goat serum (v/v) for 1hr at 37°C for antigen blocking, and then incubated overnight at 4°C with the following primary antibodies: mouse anti-tyrosine hydroxylase (TH, 1:1000, MAB318, Millipore), rabbit anti-glial fibrillary acidic protein (GFAP, 1:2000; Z033429, Dako, Denmark) or anti-Ionized calcium-binding adaptor molecule-1(Iba-1, 1:1000, 019-19741, Wako, Japan), respectively. Appropriated secondary antibodies, coupled to FITC-conjugated goat anti-mouse IgG (1:1000, A0568, Beyotime, China) or CY3-conjugated goat anti-rabbit IgG (1:1000, A0516, Beyotime, China) were applied in the dark for 1hr at 37°C to detected the primary antibodies. Finally, the tissue slices were counterstained with 4′,6-diamidino-2-phenylindole (DAPI) in mounting medium (P0131, Beyotime, China) and imaged with a fluorescence microscope (ZEISS AXIO IMAGER).

For each animal, 6 representative brain slices in each SN level were chosen to be double stained with TH and GFAP or Iba-1, respectively. The whole left and right SN area were analyzed, the TH-positive cells were countered manually by a blind experimenter. For each mouse, the expression level of positive cells for TH, GFAP and Iba-1 were obtained separately for each SN level, and the values from all levels was averaged to obtain a mean. The abovementioned brain slices were quantified for each mouse, and each group contains 6 mice.

### Hematoxylin–eosin staining and histological analysis

Colon tissues were fixed in 4% PFA and embedded in paraffin, then cut into 3 mm sections by microtome. Colon sections were deparaffinized and rehydrated by xylene and declining grades of ethanol (100, 90, 80, and 70% ethanol) for 5 min in each grade, and after 3 washes in PBS (pH 7.4) for 5 min each. For hematoxylin–eosin (HE) staining, the brain slides were stained with HE. Photomicrographs were taken with an 3D HSTECH (Hungary). Observation for histomorphological change were performed with Case Viewer (v2.4).

### Western blot analysis

Striatum and colon tissues were homogenized in ice-cold RIPA buffer (Beyotime, China) supplemented with a protease and phosphatase inhibitor cocktail (Solarbio, China), the homogenate was centrifuged at 13,000 rpm, 4°C for 10 min to collect supernatant, protein in the supernatant was determined using a BCA kit (Solarbio, China) with bovine serum albumin as a standard. The supernatant samples were denatured by boiling for 10min with loading buffer. 40μg of total proteins from each sample were separated by 8∼12% SDS-PAGE and transferred onto PVDF membranes (Millipore, USA), the membranes were blocked in 5% BSA, and probed with overnight at 4°C with primary antibodies, then incubated with HRP-conjugated secondary antibodies. Finally, the protein bands were visualized through the addition of super enhanced chemiluminescent (ECL) (Millipore, USA) for exposure and imaged by a Gel Image System (Bio-Rad, Hercules, CA, USA). The content of target protein was normalized to GAPDH or β-Actin, the densitometry was analyzed by Image J software.

Primary antibodies: mouse anti-tyrosine hydroxylase (TH, 1:1000, MAB318, Millipore), rabbit anti-BDNF (1:1000, ab108319, Abcam), rabbit anti-TLR4 (1:1000, 19811-1-AP, Proteintech), rabbit anti-NF-*κ*B (1:1000, #8242, Cell Signaling Technology), rabbit anti-MyD88(1:1000, #4283, Cell Signaling Technology), rabbit anti-occludin (13409-1-AP, Proteintech), rabbit anti-ZO-1 (1:1000, ab9658, Abcam), mouse anti-β-Actin (1:1000, 10,068-1-AP, Proteintech) and rabbit anti-GAPDH (1:8000, 60,001-1-Ig, Proteintech). Secondary antibodies: Goat anti-rabbit IgG (1:8000, B900210, Proteintech) and goat anti-mouse IgG (1:4000, 15070, China) conjugated to horseradish peroxidase.

### 16S rRNA gene sequence analyses for gut microbiome

Fresh stool samples from mice were collected and stored at −80°C. Total genomic DNA from fecal samples was extracted using a Fast DNA Spin Kit for Feces (MP Biomedical, 6570200, USA) according to the manufacturer’s instructions. The bacterial 16S rDNA V3-V4 region was amplified using barcoded 341F/806R primers (forward primer, 5′-CCTAYGGGRBGCASCAG-3′, reverse primer: 5′-GGACTACNNGGGTATCTAAT-3′) under the following PCR conditions: 95°C for 5min, followed by 30 cycles of 95°C for 30s, 55°C for 30s, 72°C for 30s, and a final extension of 72°C for 7min, then the reaction temperature was held at 12°C (Klindworth et al., 2013; Zhang et al., 2017; Zhao et al., 2013). The PCR products were purified by using a DNA Gel/PCR Purification Miniprep kit (Biomiga, BW-DC3511-01, China) and quantified with a Quant-iT PicoGreen dsDNA Assay Kit (Life Technologies, Carlsbad, USA). Libraries were prepared using TruSeq DNA LT Sample Preparation Kits (Illumina, FC-121-2001, USA). DNA sequencing was performed on Illumina Miseq platform.

16S rRNA sequence data were analyzed by the QIIME v1.9.1 platform. The raw sequences were filtered by removing short length (<200 bp) and low-quality (<30bp) sequences, only high-quality reads for bioinformatics were selected for further analysis. All of the valid reads with sequences similarity >97% were clustered into operational taxonomic units (OTUs), then compared to the Greengenes database (13.5) on the basis of OTU homology and classified as species. α-Diversity was estimated based on species richness and presented as Chao1 diversity index, β-diversity was used to evaluate the variation between the experimental groups presented by principal coordinate analysis (PCoA) plots.

## Determination of fecal SCFAs concentrations

Fecal SCFAs determination was performed on a gas chromatography-mass spectrometer (GC-MS-QP2010 Ultra system, Shimadzu Corporation, Japan) fitted with a Rtx-Wax column (30m × 0.25mm × 0.25μm). Helium was served as the carrier gas at the flow rate 2ml/min and split ration 10:1. volume of injection was 1μL. Temperature of the injector and the ionization was 240°C and 220°C, respectively. Column temperature was programmed to escalated from 100 °C to 140 °C for 3min at 7.5 °C per min, then to 200°C at 60°C per min. 50mg freeze-dried feces sample were homogenized in saturated NaCl solution and acidified with 10% sulfuric acid (v/v), the total SCFAs were extracted through the addition of diethyl ether and centrifuged at 14000rpm, 4°C for 15min, the ether layer was prepared for GC-MS analysis. Reference standards of acetic acid (71251, Sigma-Aldrich), propionic acid (94425, Sigma-Aldrich), butyric acid (19215, Sigma-Aldrich), Isobutyric acid (46935-U, Sigma-Aldrich), valeric acid (75054, Sigma-Aldrich) and Isovaleric acid (78651, Sigma-Aldrich) were mixed in diethyl ether to construct the standard curves, the content of SCFAs in each sample were calculated by standard curve method.

### Statistical analysis

Data are expressed as the mean ± SEM. Statistical analyses were conducted using GraphPad Prism Software 8.2.1. Differences were analyzed by one-way analysis of variance (ANOVA) with Tukey post hoc test for multiple groups comparisons. P values of <0.05 was set as the threshold for significance (*p <0.05, **p<0.01, ***p<0.001).

## Acknowledgements

This work was supported by the China Postdoctoral Science Foundation (2020M680064),and the National Key Research and Development Program of China (No. 2017YFC1601704).

## Author contributions

Ning Wang, designed and performed research, analysed the data and wrote the manuscript; Xin-Yue Pan advised on microbiota-related experiments; Hong-Kang Zhu contributed to the treatments for animals. Ya-Hui Guo reviewed and edited manuscript, He-Qian supervised and designed research. All authors have read and approved the final manuscript.

## Reference

Anitha, M., Vijay-Kumar, M., Sitaraman, S. V., Gewirtz, A. T., & Srinivasan, S. (2012a). Gut Microbial Products Regulate Murine Gastrointestinal Motility via Toll-Like Receptor 4 Signaling. Gastroenterology, 143(4), 1006-+. doi:10.1053/j.gastro.2012.06.034

Baizabal-Carvallo, J. F., & Alonso-Juarez, M. (2020). The Link between Gut Dysbiosis and Neuroinflammation in Parkinson’s Disease. Neuroscience, 432, 160–173. doi:https://doi.org/10.1016/j.neuroscience.2020.02.030

Barajon, I., Serrao, G., Arnaboldi, F., Opizzi, E., Ripamonti, G., Balsari, A., & Rumio, C. (2009). Toll-like Receptors 3, 4, and 7 Are Expressed in the Enteric Nervous System and Dorsal Root Ganglia. Journal of Histochemistry & Cytochemistry, 57(11), 1013–1023. doi:10.1369/jhc.2009.953539

Braniste, V., Al-Asmakh, M., Kowal, C., Anuar, F., Abbaspour, A., Toth, M., … Pettersson, S. (2014). The gut microbiota influences blood-brain barrier permeability in mice. Science Translational Medicine, 6(263). doi:ARTN263ra15810.1126/scitranslmed.3009759

Campolo, M., Paterniti, I., Siracusa, R., Filippone, A., Esposito, E., & Cuzzocrea, S. (2019). TLR4 absence reduces neuroinflammation and inflammasome activation in Parkinson’s diseases in vivo model. Brain Behav Immun, 76, 236–247. doi:10.1016/j.bbi.2018.12.003

Castano, A., Herrera, A. J., Cano, J., & Machado, A. (1998). Lipopolysaccharide intranigral injection induces inflammatory reaction and damage in nigrostriatal dopaminergic system. J Neurochem, 70(4), 1584–1592. doi:10.1046/j.1471-4159.1998.70041584.x

Cote, M., Poirier, A. A., Aube, B., Jobin, C., Lacroix, S., & Soulet, D. (2015). Partial depletion of the proinflammatory monocyte population is neuroprotective in the myenteric plexus but not in the basal ganglia in a MPTP mouse model of Parkinson’s disease. Brain Behavior and Immunity, 46, 154–167. doi:10.1016/j.bbi.2015.01.009

Cryan, J. F., O’Riordan, K. J., Cowan, C. S. M., Sandhu, K. V., Bastiaanssen, T. F. S., Boehme, M., … Dinan, T. G. (2019). The Microbiota-Gut-Brain Axis. Physiol Rev, 99(4), 1877–2013. doi:10.1152/physrev.00018.2018

Dalile, B., Van Oudenhove, L., Vervliet, B., & Verbeke, K. (2019). The role of short-chain fatty acids in microbiota-gut-brain communication. Nat Rev Gastroenterol Hepatol, 16(8), 461–478. doi:10.1038/s41575-019-0157-3

Devos, D., Lebouvier, T., Lardeux, B., Biraud, M., Rouaud, T., Pouclet, H., … Derkinderen, P. (2013). Colonic inflammation in Parkinson’s disease. Neurobiology of Disease, 50, 42–48. doi:10.1016/j.nbd.2012.09.007

Di Persio, S., Starace, D., Capponi, C., Saracino, R., Fera, S., Filippini, A., & Vicini, E. (2020). TNF-alpha inhibits GDNF levels in Sertoli cells, through a NF-kappaB-dependent, HES1-dependent mechanism. Andrology. doi:10.1111/andr.12959

Erickson, J. T., Brosenitsch, T. A., & Katz, D. M. (2001). Brain-derived neurotrophic factor and glial cell line-derived neurotrophic factor are required simultaneously for survival of dopaminergic primary sensory neurons in vivo. Journal of Neuroscience, 21(2), 581–589. doi:Doi 10.1523/Jneurosci.21-02-00581.2001

Erny, D., Hrabe de Angelis, A.L., Jaitin, D., Wieghofer, P., Staszewski, O., David, E., … Prinz, M. (2015). Host microbiota constantly control maturation and function of microglia in the CNS. Nat Neurosci, 18(7), 965–977. doi:10.1038/nn.4030

Finegold, S. M., Dowd, S. E., Gontcharova, V., Liu, C., Henley, K. E., Wolcott, R. D., … Green, J. A. (2010). Pyrosequencing study of fecal microflora of autistic and control children. Anaerobe, 16(4), 444–453. doi:https://doi.org/10.1016/j.anaerobe.2010.06.008

Gerhardt, S., & Mohajeri, M. H. (2018). Changes of Colonic Bacterial Composition in Parkinson’s Disease and Other Neurodegenerative Diseases. Nutrients, 10(6). doi:ARTN70810.3390/nu10060708

Gershanik, O. S. (2018). Does Parkinson’s disease start in the gut? Arquivos De Neuro-Psiquiatria, 76(2), 67–70. doi:10.1590/0004-282x20170188

Gorecki, A. M., Preskey, L., Bakeberg, M. C., Kenna, J. E., Gildenhuys, C., MacDougall, G., … Anderton, R. S. (2019). Altered Gut Microbiome in Parkinson’s Disease and the Influence of Lipopolysaccharide in a Human alpha-Synuclein Over-Expressing Mouse Model. Frontiers in Neuroscience, 13. doi:ARTN83910.3389/fnins.2019.00839

Grenham, S., Clarke, G., Cryan, J. F., & Dinan, T. G. (2011). Brain-gut-microbe communication in health and disease. Frontiers in Physiology, 2. doi:ARTN9410.3389/fphys.2011.00094

Hanke, M. L., & Kielian, T. (2011). Toll-like receptors in health and disease in the brain: mechanisms and therapeutic potential. Clinical Science, 121(9-10), 367–387. doi:10.1042/Cs20110164

Henke, M. T., Kenny, D. J., Cassilly, C. D., Vlamakis, H., Xavier, R. J., & Clardy, J. (2019). Ruminococcus gnavus, a member of the human gut microbiome associated with Crohn’s disease, produces an inflammatory polysaccharide. Proc Natl Acad Sci U S A, 116(26), 12672–12677. doi:10.1073/pnas.1904099116

Huh, E., Choi, J. G., Noh, D., Yoo, H. S., Ryu, J., Kim, N. J., … Oh, M. S. (2020). Ginger and 6-shogaol protect intestinal tight junction and enteric dopaminergic neurons against 1-methyl-4-phenyl 1,2,3,6-tetrahydropyridine in mice. Nutr Neurosci, 23(6), 455–464. doi:10.1080/1028415X.2018.1520477

Imamura, K., Hishikawa, N., Ono, K., Suzuki, H., Sawada, M., Nagatsu, T., … Hashizume, Y. (2005). Cytokine production of activated microglia and decrease in neurotrophic factors of neurons in the hippocampus of Lewy body disease brains. Acta Neuropathologica, 109(2), 141–150. doi:10.1007/s00401-004-0919-y

Jakobsson, H. E., Rodriguez-Pineiro, A. M., Schutte, A., Ermund, A., Boysen, P., Bemark, M., … Johansson, M. E. V. (2015). The composition of the gut microbiota shapes the colon mucus barrier. Embo Reports, 16(2), 164–177. doi:DOI 10.15252/embr.201439263

Jang, J. H., Yeom, M. J., Ahn, S., Oh, J. Y., Ji, S., Kim, T. H., & Park, H. J. (2020). Acupuncture inhibits neuroinflammation and gut microbial dysbiosis in a mouse model of Parkinson’s disease. Brain Behavior and Immunity, 89, 641–655. doi:10.1016/j.bbi.2020.08.015

Jankovic, J. (2008). Parkinson’s disease: clinical features and diagnosis. J Neurol Neurosurg Psychiatry, 79(4), 368–376. doi:10.1136/jnnp.2007.131045

Kelly, C. J., Zheng, L., Campbell, E. L., Saeedi, B., Scholz, C. C., Bayless, A. J., … Colgan, S. P. (2015). Crosstalk between Microbiota-Derived Short-Chain Fatty Acids and Intestinal Epithelial HIF Augments Tissue Barrier Function. Cell Host & Microbe, 17(5), 662–671. doi:10.1016/j.chom.2015.03.005

Kesika, P., Suganthy, N., Sivamaruthi, B. S., & Chaiyasut, C. (2021). Role of gut-brain axis, gut microbial composition, and probiotic intervention in Alzheimer’s disease. Life Sciences, 264. doi:ARTN11862710.1016/j.lfs.2020.118627

Kim, J. S., & Sung, H. Y. (2015). Gastrointestinal Autonomic Dysfunction in Patients with Parkinson’s Disease. Journal of Movement Disorders, 8(2), 76–82. doi:10.14802/jmd.15008

Klindworth, A., Pruesse, E., Schweer, T., Peplies, J., Quast, C., Horn, M., & Glockner, F. O. (2013). Evaluation of general 16S ribosomal RNA gene PCR primers for classical and next-generation sequencing-based diversity studies. Nucleic Acids Res, 41(1), e1. doi:10.1093/nar/gks808

Kuribara, H., Higuchi, Y., & Tadokoro, S. (1977). Effects of central depressants on rota-rod and traction performances in mice. Jpn J Pharmacol, 27(1), 117–126. doi:10.1254/jjp.27.117

LeBlanc, J. G., Milani, C., de Giori, G. S., Sesma, F., van Sinderen, D., & Ventura, M. (2013). Bacteria as vitamin suppliers to their host: a gut microbiota perspective. Current Opinion in Biotechnology, 24(2), 160–168. doi:10.1016/j.copbio.2012.08.005

Luchtman, D. W., Shao, D., & Song, C. (2009). Behavior, neurotransmitters and inflammation in three regimens of the MPTP mouse model of Parkinson’s disease. Physiology & Behavior, 98(1-2), 130–138. doi:10.1016/j.physbeh.2009.04.021

Morais, L. H., Hara, D. B., Bicca, M. A., Poli, A., & Takahashi, R. N. (2018). Early signs of colonic inflammation, intestinal dysfunction, and olfactory impairments in the rotenone-induced mouse model of Parkinson’s disease. Behavioural Pharmacology, 29(2-3), 199–210. doi:10.1097/Fbp.0000000000000389

Okun, E., Griffioen, K. J., Lathia, J. D., Tang, S. C., Mattson, M. P., & Arumugam, T. V. (2009). Toll-like receptors in neurodegeneration. Brain Research Reviews, 59(2), 278–292. doi:10.1016/j.brainresrev.2008.09.001

Ono, S., Karaki, S., & Kuwahara, A. (2004). Short-chain fatty acids decrease the frequency of spontaneous contractions of longitudinal muscle via enteric nerves in rat distal colon. Japanese Journal of Physiology, 54(5), 483–493. doi:10.2170/jjphysiol.54.483

Ortega-Cava, C. F., Ishihara, S., Rumi, M. A. K., Kawashima, K., Ishimura, N., Kazumori, H., … Kinoshita, Y. (2003). Strategic compartmentalization of toll-like receptor 4 in the mouse gut. Journal of Immunology, 170(8), 3977–3985. doi:10.4049/jimmunol.170.8.3977

Palasz, E., Bak, A., Gasiorowska, A., & Niewiadomska, G. (2017). The role of trophic factors and inflammatory processes in physical activity-induced neuroprotection in Parkinson’s disease. Postepy Higieny I Medycyny Doswiadczalnej, 71, 713-726. Retrieved from <Go to ISI>://WOS:000419353500001

Perez-Pardo, P., Dodiya, H. B., Engen, P. A., Forsyth, C. B., Huschens, A. M., Shaikh, M., … Keshavarzian, A. (2019). Role of TLR4 in the gut-brain axis in Parkinson’s disease: a translational study from men to mice. Gut, 68(5), 829–843. doi:10.1136/gutjnl-2018-316844

Perlmutter, J. S., & Ushe, M. (2020). Parkinson’s Disease -What’s the FUS? N Engl J Med, 383(26), 2582–2584. doi:10.1056/NEJMe2031151

Ploger, S., Stumpff, F., Penner, G. B., Schulzke, J. D., Gabel, G., Martens, H., … Aschenbach, J. R. (2012). Microbial butyrate and its role for barrier function in the gastrointestinal tract. Barriers and Channels Formed by Tight Junction Proteins Ii, 1258, 52–59. doi:10.1111/j.1749-6632.2012.06553.x

Qian, Y. W., Yang, X. D., Xu, S. Q., Wu, C. Y., Song, Y. Y., Qin, N., … Xiao, Q. (2018). Alteration of the fecal microbiota in Chinese patients with Parkinson’s disease. Brain Behavior and Immunity, 70, 194–202. doi:10.1016/j.bbi.2018.02.016

Rosser, E. C., Oleinika, K., Tonon, S., Doyle, R., Bosma, A., Carter, N. A., … Mauri, C. (2014). Regulatory B cells are induced by gut microbiota-driven interleukin-1beta and interleukin-6 production. Nat Med, 20(11), 1334–1339. doi:10.1038/nm.3680

Salat-Foix, D., Tran, K., Ranawaya, R., Meddings, J., & Suchowersky, O. (2012). Increased Intestinal Permeability and Parkinson Disease Patients: Chicken or Egg? Canadian Journal of Neurological Sciences, 39(2), 185–188. doi:10.1017/S0317167100013202

Sampson, T. R., Debelius, J. W., Thron, T., Janssen, S., Shastri, G. G., Ilhan, Z. E., … Mazmanian, S. K. (2016). Gut Microbiota Regulate Motor Deficits and Neuroinflammation in a Model of Parkinson’s Disease. Cell, 167(6), 1469-1480.e1412. doi:https://doi.org/10.1016/j.cell.2016.11.018

Sampson, T. R., Debelius, J. W., Thron, T., Janssen, S., Shastri, G. G., Ilhan, Z. E., … Mazmanian, S. K. (2016). Gut Microbiota Regulate Motor Deficits and Neuroinflammation in a Model of Parkinson’s Disease. Cell, 167(6), 1469-+. doi:10.1016/j.cell.2016.11.018

Schroeder, P., Rivalan, M., Zaqout, S., Kruger, C., Schuler, J., Long, M., … Lehnardt, S. (2021). Abnormal brain structure and behavior in MyD88-deficient mice. Brain Behavior and Immunity, 91, 181–193. doi:10.1016/j.bbi.2020.09.024

Shaw, M. H., Kamada, N., Kim, Y. G., & Nunez, G. (2012). Microbiota-induced IL-1 beta, but not IL-6, is critical for the development of steady-state T(H)17 cells in the intestine. Journal of Experimental Medicine, 209(2), 251–258. doi:10.1084/jem.20111703

Sun, M.-F., Zhu, Y.-L., Zhou, Z.-L., Jia, X.-B., Xu, Y.-D., Yang, Q., … Shen, Y.-Q. (2018). Neuroprotective effects of fecal microbiota transplantation on MPTP-induced Parkinson’s disease mice: Gut microbiota, glial reaction and TLR4/TNF-α signaling pathway. Brain, Behavior, and Immunity, 70, 48–60. doi:https://doi.org/10.1016/j.bbi.2018.02.005

Tong, L., Balazs, R., Soiampornkul, R., Thangnipon, W., & Cotman, C. W. (2008). Interleukin-1 beta impairs brain derived neurotrophic factor-induced signal transduction. Neurobiol Aging, 29(9), 1380–1393. doi:10.1016/j.neurobiolaging.2007.02.027

Topping, D. L., & Clifton, P. M. (2001). Short-chain fatty acids and human colonic function: roles of resistant starch and nonstarch polysaccharides. Physiol Rev, 81(3), 1031–1064. doi:10.1152/physrev.2001.81.3.1031

van Noort, J. M., & Bsibsi, M. (2009). Toll-like receptors in the CNS: implications for neurodegeneration and repair. Neurotherapy: Progress in Restorative Neuroscience and Neurology, 175, 139–148. doi:10.1016/S0079-6123(09)17509-X

Verbaan, D., Marinus, J., Visser, M., van Rooden, S. M., Stiggelbout, A. M., & van Hilten, J. J. (2007). Patient-reported autonomic symptoms in Parkinson disease. Neurology, 69(4), 333–341. doi:10.1212/01.wnl.0000266593.50534.e8

Verma, R., Cronin, C. G., Hudobenko, J., Venna, V. R., McCullough, L. D., & Liang, B. T. (2017). Deletion of the P2X4 receptor is neuroprotective acutely, but induces a depressive phenotype during recovery from ischemic stroke. Brain Behavior and Immunity, 66, 302–312. doi:10.1016/j.bbi.2017.07.155

Villaran, R. F., Espinosa-Oliva, A. M., Sarmiento, M., de Pablos, R. M., Arguelles, S., Delgado-Cortes, M. J., … Machado, A. (2010). Ulcerative colitis exacerbates lipopolysaccharide-induced damage to the nigral dopaminergic system: potential risk factor in Parkinson’s disease. Journal of Neurochemistry, 114(6), 1687–1700. doi:10.1111/j.1471-4159.2010.06879.x

Vinolo, M. A. R., Rodrigues, H. G., Nachbar, R. T., & Curi, R. (2011). Regulation of Inflammation by Short Chain Fatty Acids. Nutrients, 3(10), 858–876. doi:10.3390/nu3100858

Vogt, N. M., Kerby, R. L., Dill-McFarland, K. A., Harding, S. J., Merluzzi, A. P., Johnson, S. C., … Rey, F. E. (2017). Gut microbiome alterations in Alzheimer’s disease. Scientific Reports, 7(1), 13537. doi:10.1038/s41598-017-13601-y

Westfall, S., Lomis, N., Kahouli, I., Dia, S. Y., Singh, S. P., & Prakash, S. (2017). Microbiome, probiotics and neurodegenerative diseases: deciphering the gut brain axis. Cellular and Molecular Life Sciences, 74(20), 3769–3787. doi:10.1007/s00018-017-2550-9

Xiong, M. G., Xu, Z. S., Li, Y. H., Wang, S. Y., Wang, Y. Y., & Ran, Y. (2020). RNF152 positively regulates TLR/IL-1R signaling by enhancing MyD88 oligomerization. Embo Reports, 21(3). doi:ARTNe4886010.15252/embr.201948860

Xu, H., Wang, Y., Song, N., Wang, J., Jiang, H., & Xie, J. (2017). New Progress on the Role of Glia in Iron Metabolism and Iron-Induced Degeneration of Dopamine Neurons in Parkinson’s Disease. Front Mol Neurosci, 10, 455. doi:10.3389/fnmol.2017.00455

Zhang, J., Zhu, C., Guan, R., Xiong, Z., Zhang, W., Shi, J., … Lu, Z. (2017). Microbial profiles of a drinking water resource based on different 16S rRNA V regions during a heavy cyanobacterial bloom in Lake Taihu, China. Environ Sci Pollut Res Int, 24(14), 12796–12808. doi:10.1007/s11356-017-8693-2

Zhao, L., Wang, G., Siegel, P., He, C., Wang, H., Zhao, W., … Meng, H. (2013). Quantitative genetic background of the host influences gut microbiomes in chickens. Sci Rep, 3, 1163. doi:10.1038/srep01163

Zhou, Z. L., Jia, X. B., Sun, M. F., Zhu, Y. L., Qiao, C. M., Zhan, B. P., … Shen, Y. Q. (2019). Neuroprotection of Fasting Mimicking Diet on MPTP-Induced Parkinson’s Disease Mice via Gut Microbiota and Metabolites. Neurotherapeutics, 16(3), 741–760. doi:10.1007/s13311-019-00719-2

